# Bacterial genome reduction as a result of short read sequence assembly

**DOI:** 10.1101/091314

**Authors:** Charles H.D. Williamson, Andrew Sanchez, Adam Vazquez, Joshua Gutman, Jason W. Sahl

## Abstract

High-throughput comparative genomics has changed our view of bacterial evolution and relatedness. Many genomic comparisons, especially those regarding the accessory genome that is variably conserved across strains in a species, are performed using assembled genomes. For completed genomes, an assumption is made that the entire genome was incorporated into the genome assembly, while for draft assemblies, often constructed from short sequence reads, an assumption is made that genome assembly is an approximation of the entire genome. To understand the potential effects of short read assemblies on the estimation of the complete genome, we downloaded all completed bacterial genomes from GenBank, simulated short reads, assembled the simulated short reads and compared the resulting assembly to the completed assembly. Although most simulated assemblies demonstrated little reduction, others were reduced by as much as 25%, which was correlated with the repeat structure of the genome. A comparative analysis of lost coding region sequences demonstrated that up to 48 CDSs or up to ~112,000 bases of coding region sequence, were missing from some draft assemblies compared to their finished counterparts. Although this effect was observed to some extent in 32% of genomes, only minimal effects were observed on pan-genome statistics when using simulated draft genome assemblies. The benefits and limitations of using draft genome assemblies should be fully realized before interpreting data from assembly-based comparative analyses.

## Introduction

Advances in DNA sequencing technologies have allowed for large-scale whole genome sequencing of bacterial genomes. Short read technologies, such as those employed on the Illumina sequencing platforms, have facilitated high-throughput analyses of organisms for the purposes of comparative genomics (1), phylogeography (2), and association of genomic attributes with antimicrobial resistance (3). While reference-guided methods, including the identification of single nucleotide polymorphisms (SNPs), are important for understanding population genetics (4), many analyses are typically performed with assembled genomes making genome assembly an important and standard method in the analysis of bacterial organisms.

Studies that rely on assembled genomes include analyzing the conservation of genomic features within a set of isolates and estimating core and pan-genomes. Core and pan-genome analyses, introduced by Tettelin and colleagues (5), have been applied to many bacterial species (6), and a number of tools have been developed to calculate and analyze the pan-genome (7–13). All of these tools rely on assembled genomes (or protein/nucleotide sequences from assemblies) as input. Most of the assembled genomes currently available in public databases are draft assemblies. Of the approximately 80,000 bacterial genomes available from NCBI on November 1, 2016, less than 6000 are complete.

As assemblies generated from short read sequencing data have become an integral part of many research projects, potential limitations of this type of data must be considered. For instance, contaminating reads can be incorporated into assemblies (14–16) requiring post-assembly screening and quality control. Additionally, genome assemblies generated from short read technologies are typically fragmented due to the inability of short reads (and insert regions) to span large repeat regions of a genome (17), which often breaks assemblies into multiple contigs. This fragmentation can drop genomic regions from an assembly, which look like missing regions in comparative analyses. In this study, we evaluated how well assemblies generated from short read data estimate complete bacterial genomes.

## Methods and Materials

**Complete Genomes Used**. We downloaded (September 16, 2016) all bacterial genomes from GenBank, then filtered the genomes to only include completed assemblies (n=5676). We then filtered out genomes that contained >10 non-nucleotide characters (non A,T,G,C), which could indicate problems with genome assembly (n=203). A complete list of genomes (n=5473) used in this study is shown in Table S1.

**Read simulation**. Paired end illumina reads were simulated for each complete genome using ART (18) vMountRainier with the following parameters: -ss MSv3 - l 250 -f 75 -m 300 -s 30. Genomes were then assembled with SPAdes v3.7.1 (19) using the following parameters: -t 4 -k 21,33,55,77,99,127 -cov-cutoff auto - careful -1 pair1 -2 pair2. Following assembly, genomes were polished with Pilon 2 v.1.7, using the following parameters: -threads 4 -fix all,amb. Contigs shorter than 200bp were filtered from the assembly to stay consistent with GenBank standards. The genome assembly was automated with the UGAP assembly pipeline (https://github.com/jasonsahl/UGAP), which was run using the Slurm management system on a high-performance computing (HPC) cluster at Northern Arizona University.

In order to identify how well the simulated reads represented the completed genomes, we mapped the reads to the completed genome with BWA-MEM (20). The per base coverage was calculated with the GenomeCoverageBed method in BEDTOOLS (21). The number of bases with a minimum coverage of 1 was then divided by the total number of bases in the completed genome to calculate the percent coverage of simulated reads across each genome.

**Genome validation**. In addition to simulated reads, we also analyzed a set of 49 complete, or near complete, genomes that have been assembled separately with both Illumina and PacBio sequencing platforms (Table S2). To test the ability of ART to simulate representative short sequencing reads, we ran the Illumina reads through SPAdes using the same parameters as with the simulated reads.

**Genome size calculation.** For each genome, we summed the entire sequence length across all sequences with a Python script (https://gist.github.com/jasonsahl/64d88d2858a915ee730b5f86e305e5d4). We divided the size of the simulated assembly by the size of the completed assembly to determine the amount of the genome retained.

**Repeat characterization**. To identify the percentage of the genome associated with repeated regions, we aligned each genome against itself with NUCmer (22). We then divided the number of bases in repeated regions by the total length of the genome to characterize the repeat percentage. The identification of repeat regions was facilitated by methods implemented in the NASP pipeline (4). Using default parameters, NUCmer is unable to detect repeats shorter than 21 nucleotides.

**Multi-locus Sequence Typing comparisons**. The sequence type of *E. coli* and *S. aureus* assemblies was identified using the PubMLST system and a custom script (https://github.com/jasonsahl/mlst_blast.git). Each allele was assigned if an exact match to the database was observed.

**Comparative genomics.** To identify the impact of regions collapsed or lost during the genome assembly using simulated reads, a large-scale Blast Score Ratio (LS-BSR) (12, 23) analysis was performed. Coding regions were predicted from the completed genome and the simulated genome with Prodigal (24). All coding regions were clustered with USEARCH (25) at an ID of 0.9 and aligned against both genomes with BLAT (26). The BSR values were then compared between the simulated and the completed genome to identify the number and combined length of regions that had a BSR value > 0.8 (~80% peptide identity over 100% of the peptide length) in the completed genome and a BSR value < 4 0.4 in the simulated genome. These regions represent those that were lost from the assembly and could confound comparative analysis using genome assemblies of short read sequence data.

**Publicly available genomes**. To characterize the quality of all genomes from a single species in public databases, all *Escherichia coli* genome assemblies (n=4842) were downloaded on September 16, 2016. Genomes were assessed for contig number and assembly size.

## Results

**Extent of genome reduction using simulated, short read assemblies**. In order to understand the effects of short read assembly on the retention of sequence from bacterial genomes, we downloaded all completed genomes from GenBank with fewer than 10 ambiguities (n=5473) (Table S1) and simulated paired-end Illumina MiSeq reads with ART (18) at an average coverage of 75x. We assembled all genomes with SPAdes as it performs well compared to other assemblers (27), it recovers larger portions of reference genomes than other short read assembly algorithms (28), and we wanted to keep the assembly algorithm constant. The sizes of the complete and the simulated genomes were compared to understand the extent of reduction due to assembly problems. While the vast majority of the genome was recovered in most cases, some genomes showed significant reduction due to short read assembly (Figure 1, Table S3). The maximum percentage of observed genome reduction was approximately 25% in *Orientia tsutsugamushi,* which has been described as having one of the most duplicated genomes (29). In some cases, the simulated genome assembly was slightly larger than the complete genome (maximum of ~0.76% larger), which may be due to the presence of contigs in the simulated genome that should have been merged during assembly.

**Figure 1:**
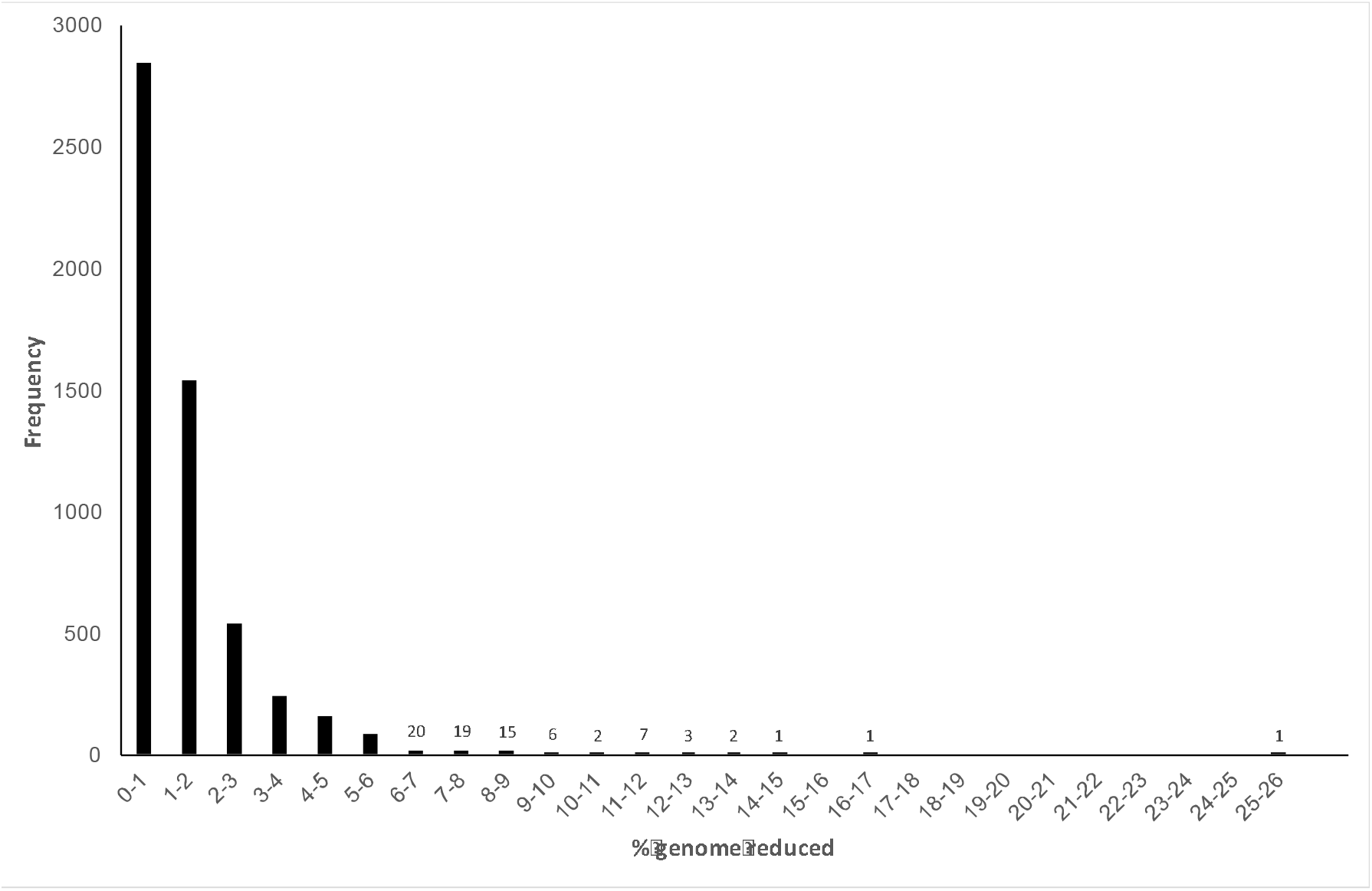
Frequency plot of the number of genomes against the extent of genome reduction.

We then calculated the breadth of coverage of the completed genome, at a minimum depth of 1x, with simulated reads (Table S3). The breadth of coverage was meant to estimate how well the simulated reads represented the complete genome. Breadth of coverage values range from approximately 73% to 100%. A correlation of breadth of coverage and genome reduction (correlation coefficient=0.76) demonstrates that different methods (genome assembly and short read mapping) return a similar result (Figure 2, Table S3).

**Figure 2:**
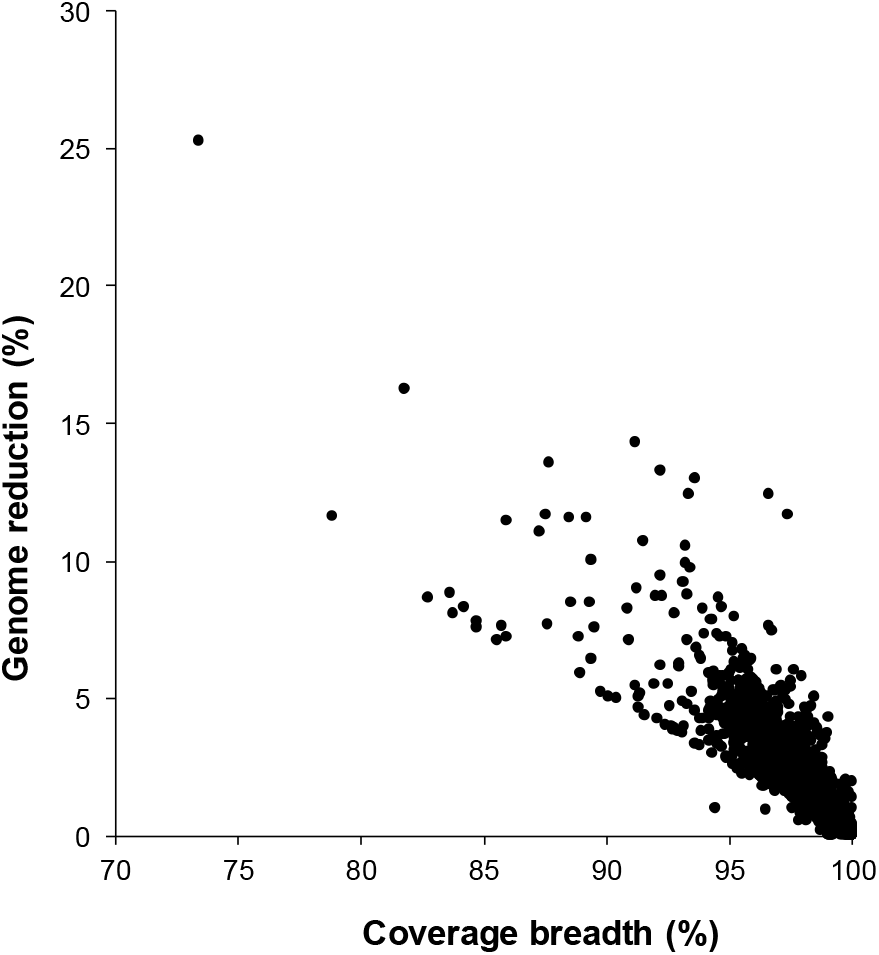
Breadth of coverage of simulated sequencing reads across complete genomes compared to the genome reduction of simulated genome assemblies compared to completed genomes.

**Genome reduction using actual sequence data**. To confirm that genome reduction wasn‘t solely due to the short read simulation, a set of 49 complete or near complete *Burkholderia* genomes (30) was compared to the same isolates where the genomes were also sequenced on the Illumina MiSeq platform. When the genome reduction percentages were compared between real and simulated reads, similar results were observed (correlation coefficient=0.50) (Table 1). In some cases, the Illumina assembly was larger than the completed genome, which may be due to bleed over between multiplexed samples on the same sequencing run (31) or assembly error. This analysis demonstrates that the simulated short reads should be generally representative of the extent of genome reduction across other species.

**Table 1:** Correlations between simulated and true assemblies

| Strain | PacBio<br>Assembly Size | Illumina<br>Assembly Size | Simulated<br>Assembly Size | %reduction<br>(Illumina) | %reduction<br>(simulated) |
| --- | --- | --- | --- | --- | --- |
| Bp7003 | 6590904 | 6498086 | 6511321 | 1.41 | 1.21 |
| Bp7004 | 7357522 | 7251364 | 7263334 | 1.44 | 1.28 |
| Bp7005 | 7159534 | 7122062 | 7120222 | 0.52 | 0.55 |
| Bp7006 | 6858431 | 6782684 | 6786320 | 1.10 | 1.05 |
| Bp7046 | 6708057 | 6592936 | 6610712 | 1.72 | 1.45 |
| Bp7047 | 6739513 | 6624496 | 6646837 | 1.71 | 1.38 |
| Bp7049 | 6785295 | 6673578 | 6683615 | 1.65 | 1.50 |
| Bp7097 | 6899618 | 6819885 | 6815275 | 1.16 | 1.22 |
| Bp7270 | 6787216 | 6701472 | 6651437 | 1.26 | 2.00 |
| Bp0071 | 7319559 | 7263748 | 7209107 | 0.76 | 1.51 |
| Bp0072 | 7135010 | 7072535 | 7077251 | 0.88 | 0.81 |
| Bp0073 | 6729272 | 6633203 | 6642243 | 1.43 | 1.29 |
| Bp0150 | 6857828 | 6807568 | 6812424 | 0.73 | 0.66 |
| Bp0379 | 6902331 | 6857306 | 6861916 | 0.65 | 0.59 |
| Bp0422 | 7961561 | 7902985 | 7909489 | 0.74 | 0.65 |
| Bp0713 | 7079942 | 7036491 | 7043175 | 0.61 | 0.52 |
| Bp2202 | 7337349 | 7298664 | 7307467 | 0.53 | 0.41 |
| Bp3330 | 7956735 | 7871833 | 7878640 | 1.07 | 0.98 |
| Bp6994 | 7273038 | 7209301 | 7216862 | 0.88 | 0.77 |
| Bp6997 | 7129510 | 7024776 | 7036295 | 1.47 | 1.31 |
| Bp7014 | 8170759 | 8126665 | 8127581 | 0.54 | 0.53 |
| Bp7021 | 6815241 | 6740025 | 6746707 | 1.10 | 1.01 |
| Bp7030 | 6330734 | 6503504 | 6304216 | -2.73 | 0.42 |
| Bp7031 | 7648890 | 7608554 | 7612789 | 0.53 | 0.47 |
| Bp7035 | 7424194 | 7378871 | 7381392 | 0.61 | 0.58 |
| Bp7043 | 8354771 | 8296564 | 8294334 | 0.70 | 0.72 |
| Bp7052 | 6767931 | 6704757 | 6702085 | 0.93 | 0.97 |
| Bp7053 | 6970128 | 6778809 | 6751479 | 2.74 | 3.14 |
| Bp7055 | 6950257 | 6771434 | 6750755 | 2.57 | 2.87 |
| Bp7062 | 7235050 | 7149035 | 7147490 | 1.19 | 1.21 |
| Bp7080 | 7250560 | 6938686 | 7161963 | 4.30 | 1.22 |
| Bp7344 | 6620927 | 6532626 | 6523523 | 1.33 | 1.47 |
| Bp7345 | 6835061 | 6735080 | 6747730 | 1.46 | 1.28 |
| Bp7347 | 6614472 | 6526402 | 6536593 | 1.33 | 1.18 |
| Bp7353 | 6764528 | 6692224 | 6699728 | 1.07 | 0.96 |
| Bp7422 | 8000745 | 7624755 | 7686199 | 4.70 | 3.93 |
| Bp7432 | 8028759 | 8096860 | 7956373 | -0.85 | 0.90 |
| Bp7434 | 8028759 | 7906481 | 7956220 | 1.52 | 0.90 |
| Bp7583 | 6883100 | 6810027 | 6817056 | 1.06 | 0.96 |
| Bp7621 | 7780595 | 7705087 | 7719455 | 0.97 | 0.79 |
| Bp7630 | 6891707 | 6797470 | 6822127 | 1.37 | 1.01 |
| Bp7634 | 6913771 | 6822861 | 6839604 | 1.31 | 1.07 |
| Bp7657 | 7583794 | 7518484 | 7536053 | 0.86 | 0.63 |
| Bp7702 | 7463455 | 7418001 | 7428963 | 0.61 | 0.46 |
| Bp7709 | 6845356 | 6764904 | 6787311 | 1.18 | 0.85 |
| Bp7064 | 7427587 | 7302761 | 7286670 | 1.68 | 1.90 |
| Bp7071 | 6784408 | 7273481 | 6721097 | -7.21 | 0.93 |

**Repeat structure of all genomes**. In order to understand the repeat structure of each genome, NUCmer self-alignments were performed on all genomes and the summed repeat regions were divided by the entire genome length. The results demonstrate that several of the genomes with a high level of reduction were also highly repetitive (Figure 3). In general, genomes with a low level of repeats also had a low level of reduction. The inability to span repeats largely explains the reduction in genome size following genome assembly. As mentioned above, genome reduction is correlated to breadth of coverage (short read mapping), which highlights the limitations of short reads in resolving repeats using independent approaches.

**Figure 3:**
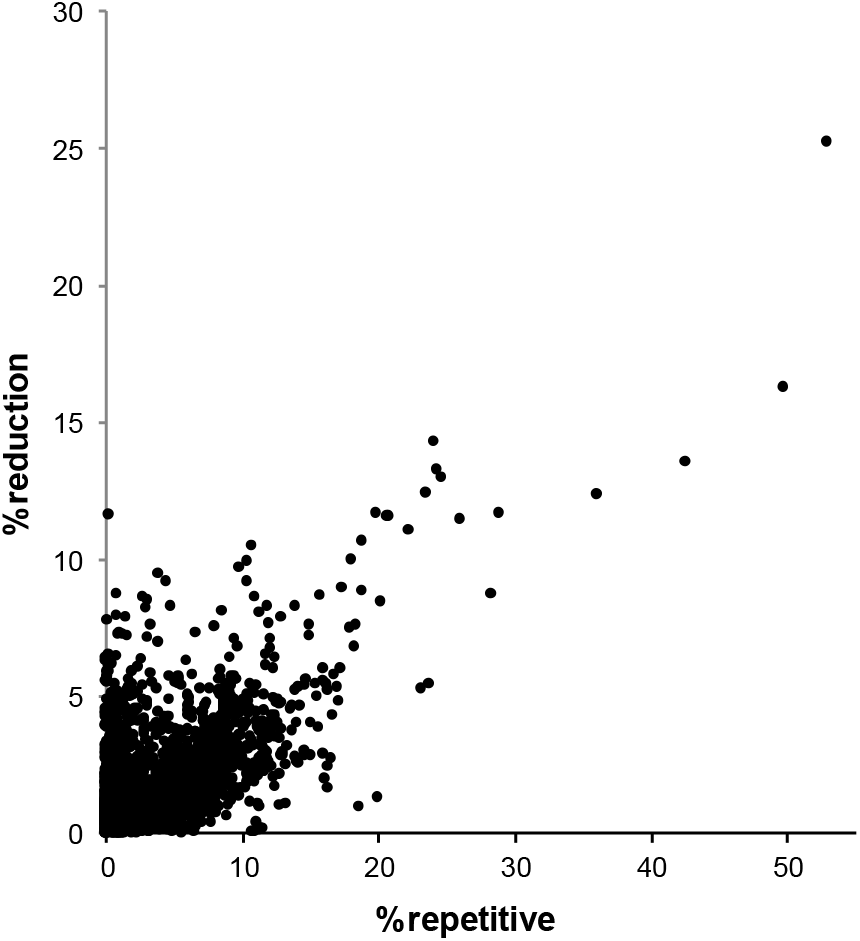
A plot of the % of the genome that is repetitive against the % of the genome that is reduced

**Draft genome assembly effects on comparative genomics**. The potential effects of a reduced genome on comparative genomics was investigated using LS-BSR. The number and length of regions that were missing from the simulated genome was calculated (Table S3). In 3729 of 5473 queried genomes, there were no coding regions (CDSs) that were missing from the simulated genome compared to the completed genome, despite seeing simulated genome assembly sizes that were up to 16% smaller than the completed genome. Of all simulated genomes, 780 were missing more than one CDS identified in the complete genome. The maximum number of CDSs missing from a simulated genome compared to a completed genome was 48, while the maximum length of coding region sequence lost in any genome was approximately 112,000 nucleotides.

Reads were aligned to CDSs identified in complete genome assemblies but missing from simulated genome assemblies to determine if short read alignment could be used to verify the presence or absence of CDSs in a genome. The breadth of coverage was determined at a minimum depth of 1X as described above. In two test cases (GCA_000017805.1, missing 48 CDSs in the simulated genome; GCA_000147815.3, missing 8 CDSs corresponding to ~112,000 nucleotides), all missing CDSs were at least partially covered by simulated reads (minimum of ~46% coverage breadth). Mapped reads provided 100% breadth of coverage for 50 of the 56 CDSs evaluated for both genomes, which suggests that read mapping is a valuable method for confirming the presence/absence of potentially missing genomic features.

**Draft genome assembly effects on pan-genome calculations.** The effect of genome reduction on core and pan genome calculations was identified in an analysis of *Escherichia coli*, *Staphylococcus aureus*, and *Salmonella enterica*, species for which numerous (>100) complete genomes are available. In each case, the core genome was calculated with LS-BSR for coding region sequences with a BSR value of > 0.8 across all genomes tested; in each case, the average core genome was calculated across 10 replicates at each level of sampling. The core genome results demonstrate the simulated and completed genomes generally return a consistent core genome size (Figure 4). Additionally, the pan-genome size was slightly larger using simulated reads, which is likely a result of fragmented coding regions that appear to be separate sequences during the clustering step in LS-BSR. The same general trends were observed across each species.

**Figure 4:**
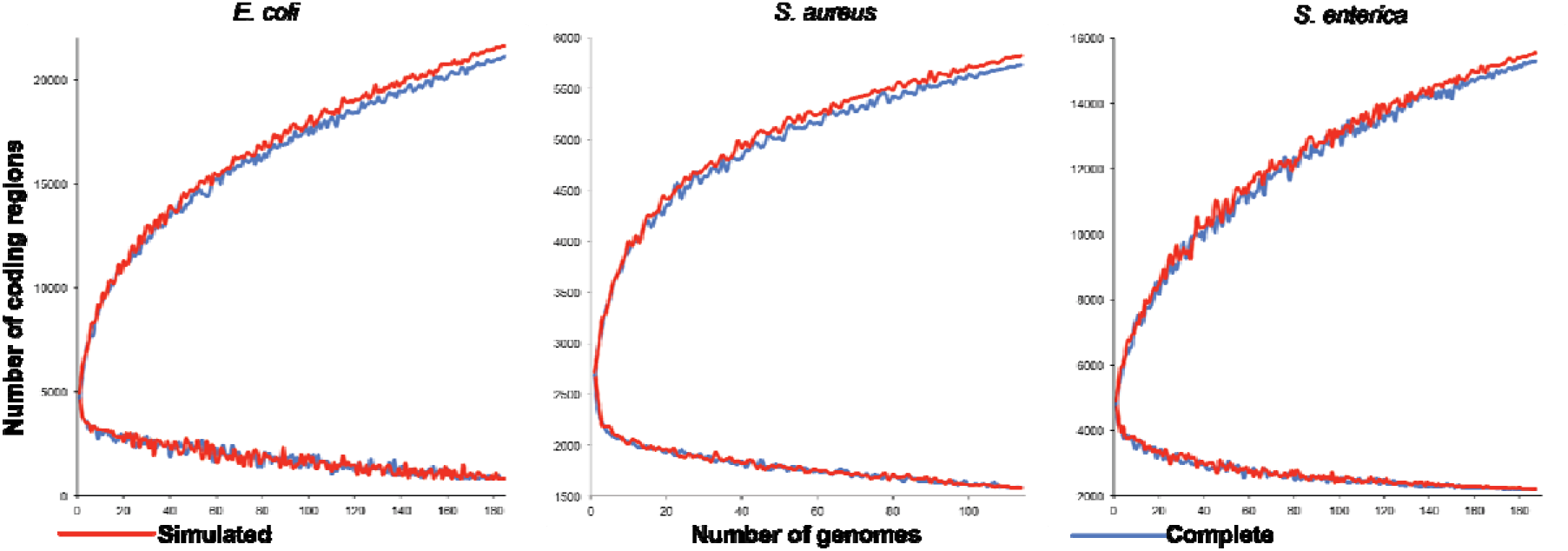
Comparative pan-genome plots for species with a large number of complete genomes. The plots either demonstrate the accumulation of coding regions in the pan-genome (upper lines) or reduction of coding regions in the core genome (lower lines).

**MLST comparisons between complete and simulated genome assemblies**. The relationships between bacterial isolates has typically been performed with multi-locus sequence type (MLST) approaches (32). To test the quality of assembled genomes, we extracted the 7 genes from the *E. coli* and *S. aureus* MLST schemes (33) and compared the sequence type (ST) calls between finished and simulated genome assemblies. In both species, all called sequence types matched between complete and simulated genome assemblies. This demonstrates that high quality draft genome assemblies can often provide important sequence type information for comparison to previous or future studies.

**Comparison of real and simulated data**. Although simulated draft genome assemblies provide comparable MLST and core genome information, they don‘t represent real data, which can be of variable quality. A comparison of contig numbers between *E. coli* genomes downloaded from GenBank and simulated assemblies generated in this study demonstrates this variability (Figure 5, panel A). The genome size is also highly variable in the real data (Figure 5, panel B), which could be due to either insufficient coverage or contamination with other genomes. If strict filtering on real genome sequence data is implemented, then much of this variation can and should be eliminated prior to comparative analyses.

**Figure 5:**
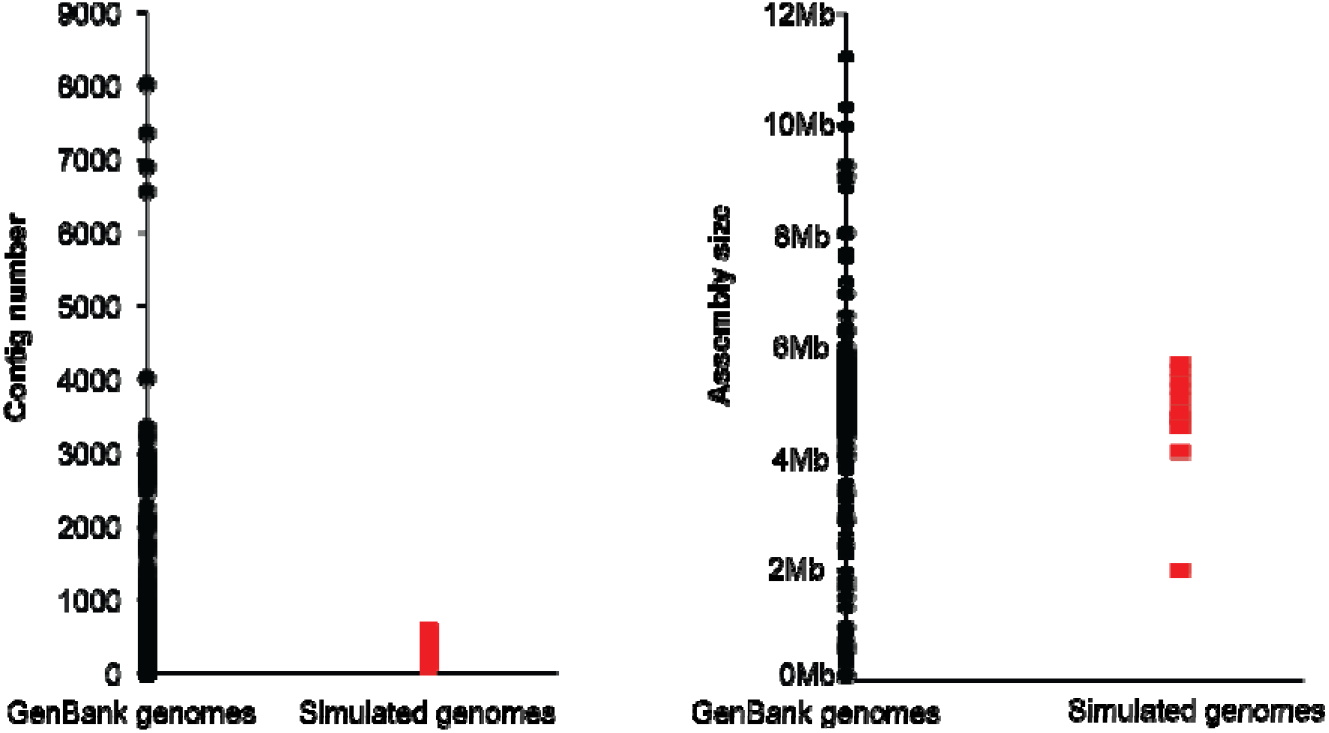
A comparison between all *Escherichia coli* genomes in Genbank (black) and simulated short read assemblies (red) in terms of (A) the number of total contigs, and (B) the summed genome assembly size.

## Discussion

Short read sequencing technologies have been key in understanding the movement (34) and population structure (35) of bacterial species. Recent advances in DNA sequencing now allow for the push button assembly of bacterial genomes using long read sequencing approaches (36), which holds the promise of automated and complete genome assembly even for highly duplicated and repetitive genomes. However, due to cost limitations, many laboratories still rely on short read technologies for high-throughput SNP identification and genome assembly ensuring that short read applications will continue to be used for large-scale comparative genomics. While benefits to these approaches exist in the consensus calling of variants, limitations also exist due to the short nature of the read composition, which depending on the length of the read and length of the repeat, cannot span many repeat regions resulting in fragmented genome assemblies. Previous work has demonstrated the effects of different genome assembly algorithms on the recovery of a reference genome using short read technologies (27, 37, 38), and the GAGE-B study (27) evaluated assemblies of eight different bacterial species genomes with a number of different assemblers.

However, less is known about how the composition of genomes from diverse species affects the ability to resolve the full genome with short read sequencing technology by keeping the assembly algorithm constant. In this study, we performed a comprehensive analysis of this issue through the assembly of greater than 5000 finished bacterial genomes. The results demonstrate that simulated short read assemblies recovered high percentages of most genomes; however, significant genome reduction was observed in some highly repetitive genomes, which has the ability to affect downstream comparative analyses.

Comparative genomics studies include the identification of genomic features that are differentially conserved between genomes from isolates in the same or closely-related species. These comparisons are important for identifying gene differences that may be associated with diagnostics, virulence, or differential phenotypes (39–44). Artifacts generated from the assembly of short read data could potentially impact these sorts of comparisons. Our results indicate that coding region sequences identified in simulated draft genome assemblies were representative of the coding regions identified in complete genomes in most cases. Thus, draft assemblies can provide important information on genomic feature variation between strains, core and pan-genome comparisons, and isolate relationships based upon MLST genes extracted from draft assembles.

The results of this study also demonstrate that draft genome quality in public repositories is variable and that quality control and filtering should be applied prior to comparative genomics studies. The results also indicate that genome reduction due to short read assembly can be a problem in downstream analyses for some genomes, although the impacts are variable, and perhaps predictable based on the repeat structure of a given genome. For large-scale comparative analyses, results must be interpreted with these limitations in mind. If missing genes are observed between groups of genomes, raw read mapping can be used to verify the gene presence or absence, although short read mapping may also suffer from some of the same limitations as short read genome assembly. Additionally, complete genomes representing species or clades of interest can provide a reference point for evaluating draft genome assemblies (e.g. provide information about repeat structure). This study indicates that draft genome assemblies generated from short read data often provide an acceptable representation of a bacterial genome for many comparative genomics applications.

## Acknowledgments

This work was facilitated by the Monsoon High Performance Cluster (HPC) resource at Northern Arizona University.

